# Depressive status modulates hippocampal-cortical dynamics during acute nicotine use

**DOI:** 10.64898/2026.03.31.715638

**Authors:** Jihye Ryu, Lisette Torres, Michael Ward, Uros Topalovic, Mauricio Vallejo Martelo, Humza Zubair, Ausaf Bari

**Affiliations:** Department of Neurosurgery, David Geffen School of Medicine at UCLA, Los Angeles, CA 90095 USA; David Geffen School of Medicine at UCLA, Los Angeles, CA 90095 USA; Department of Psychiatry and Biobehavioral Sciences, Jane and Terry Semel Institute for Neuroscience and Human Behavior, UCLA, Los Angeles, California, USA; Medical Scientist Training Program, David Geffen School of Medicine at UCLA, Los Angeles, California, USA

## Abstract

Nicotine use disorder shows heterogeneity in treatment response, potentially reflecting differences in underlying neural circuitry, particularly in the presence of depression. We examined real-time neural dynamics during nicotine inhalation in two chronic users - one with depression and one without - using simultaneous hippocampal recordings from responsive neurostimulation (RNS) electrodes and scalp EEG. Oscillatory activity and hippocampal-cortical connectivity were analyzed in relation to mood and craving. Oscillatory activity tracked mood in the non-depressed individual but was attenuated or reversed in the depressed individual, suggesting reduced reward-related neural responsiveness. In contrast, both participants showed reduced alpha hippocampal-cortical connectivity following nicotine use, suggesting a shift from reward-seeking to reward and relief processing. These findings support a network-based framework of nicotine-driven neural dynamics and provide preliminary evidence that depressive status may modulate these processes. Although limited to two cases, this work highlights the potential for identifying neurophysiological subtypes of nicotine users and informs future efforts toward personalized treatment approaches.

## Introduction

Nicotine use disorder remains a leading cause of preventable disease worldwide, contributing to cardiovascular disease, cancer, and respiratory illness [1]. Despite available pharmacological (e.g., nicotine replacement therapy, bupropion) and behavioral interventions (e.g., cognitive behavioral therapy, motivational interviewing) [2], relapse rates remain high, with many individuals returning to smoking within the first year of quitting [3]. These persistent relapses highlight substantial heterogeneity in treatment response, suggesting that current approaches fail to account for individual differences in underlying neural circuitry and reinforcing the need for more personalized interventions.

Nicotine use disorder can be conceptualized as a cyclical process involving binge/intoxication, withdrawal/negative affect, and preoccupation/anticipation, with repeated use shifting behavior from reward-driven to relief-driven [4]. However, the extent to which these processes compare across individuals remains unclear. In particular, nicotine use disorder is highly heterogeneous, and psychiatric conditions such as depression may alter this cycle by elevating baseline negative affect and reshaping the neural dynamics that sustain use [5,6].

Individuals with depression are at increased risk of nicotine dependence and exhibit altered nicotine experiences, including reduced reward sensitivity, and more persistent craving compared with those without depression [7–9]. This pattern suggests a shift toward relief-driven use, in which nicotine primarily serves to alleviate negative affect. Such differences point to variation in the neural mechanisms underlying intoxication and affect, highlighting the need to characterize these processes to support more personalized interventions.

To examine these mechanisms, we studied two chronic nicotine users-one with diagnosed depression and one without-who were implanted with responsive neurostimulation (RNS) electrodes in the hippocampus for epilepsy treatment (Figure 1A). This setup enabled direct recording of hippocampal activity in humans, which is otherwise difficult to obtain. We analyzed how hippocampal activity interacts with large-scale cortical dynamics reflected in midline EEG channels (Fz, Cz, Pz), regions implicated in craving and affective regulation [10–12]. Specifically, we examined how oscillatory activity and limbic-cortical coupling evolve during acute nicotine use in real time, and whether these dynamics differ as a function of depressive status.

**Figure 1.**
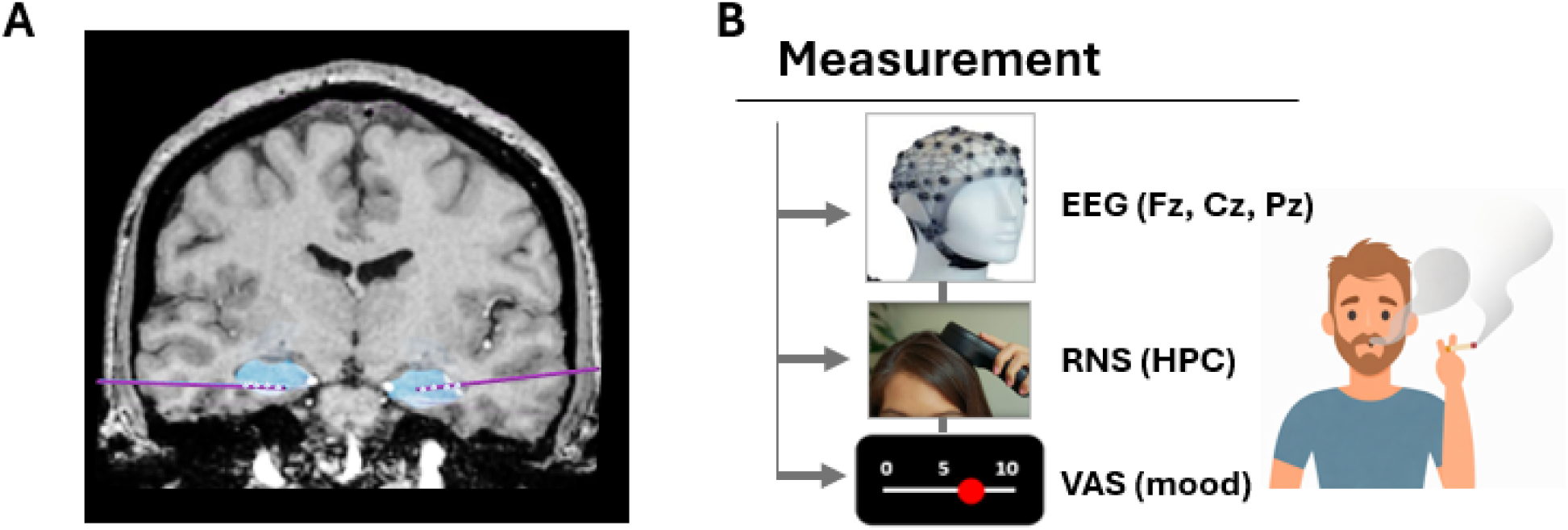
(A) Representative patient with RNS electrode leads targeting the hippocampus. (B) Experimental paradigm showing simultaneous EEG and RNS recordings alongside visual analog scale (VAS) mood ratings. HPC = hippocampus; VAS = visual analog scale.

We hypothesized that acute nicotine use would differentially modulate hippocampal-cortical dynamics as a function of depressive status, reflecting altered reward and relief processing. Specifically, we expected nicotine-related changes in functional connectivity to occur in both individuals but to be more pronounced in the participant with depression.

We further examined whether changes in oscillatory activity and hippocampal-cortical coupling tracked subjective craving during nicotine exposure.

## Methods

### Participants

Two participants with pharmaco-resistant focal epilepsy and chronic nicotine use participated in this study. Both had been implanted with a NeuroPace RNS System (Mountain View, CA) for treatment of epilepsy. Electrode placement was determined solely on the basis of clinical treatment criteria [13]. Both participants volunteered for the study and provided informed consent in accordance with a protocol approved by the UCLA Medical Institutional Review Board (IRB). Details of participant demographics and relevant clinical information are provided in Table 1.

**Table 1.** Participant Information.

| Participant ID | P1 | P2 |
| --- | --- | --- |
| Age | 54 | 26 |
| Sex | F | M |
| Current Psychiatric Diagnosis | none | depression, anxiety |
| Past Psychiatric Diagnosis | anxiety, post-traumatic stress disorder (PTSD) | depression, anxiety |
| FTND | 4 (low to moderate) | 2 (low dependence) |
| Electrode Model<sup>1</sup> | RNS 330-10 (R)<br>RNS 344-3.5 (L) | RNS 330-3.5 (R)<br>RNS 344-3.5 (L) |

| **Hemisphere** | Left, Right | Right<sup>2</sup> |
| --- | --- | --- |
| **Localization** | Hippocampus | Hippocampus |

<sup>1</sup> RNS 330-10 model had 10mm contact spacing, and others had 3.5mm contact spacing. <sup>2</sup> P2 had bilateral electrodes implanted, but only the right electrode was analyzed due to excessive noise in the left. FTND = Fagerström Test for Nicotine Dependence.

### Nicotine use task

On the day of recording, participants were enrolled in a separate study not analyzed here. During breaks between those studies, they took a smoking break while neural recordings continued. Before the break, they were asked to complete a 10-item Questionnaire of Smoking Urges (QSU) [14] and the Fagerstrom Test for Nicotine Dependence (FTND) [15]. Then, they were asked to continuously monitor and report changes in their mood while using their preferred form of nicotine (P1 smoked cigarettes; P2 vaped). Mood was measured using a visual analog scale (VAS; 0-10), where 0 indicated “sad,” 5 “neutral,” and 10 “happy.” Ratings were collected continuously using a MATLAB-based sliding scale and sampled at 60 Hz. During the task, the experimenter recorded the timing of nicotine inhalation and any other observable activities. Specifically, the first participant (P1) smoked for approximately 3 minutes and drank coffee between cigarette puffs. The second participant (P2) vaped for approximately 2 minutes and laughed naturally during the recording period. The continuously recorded mood ratings are shown in Figure 2. Following smoking/vaping, participants again completed the QSU.

**Figure 2.**
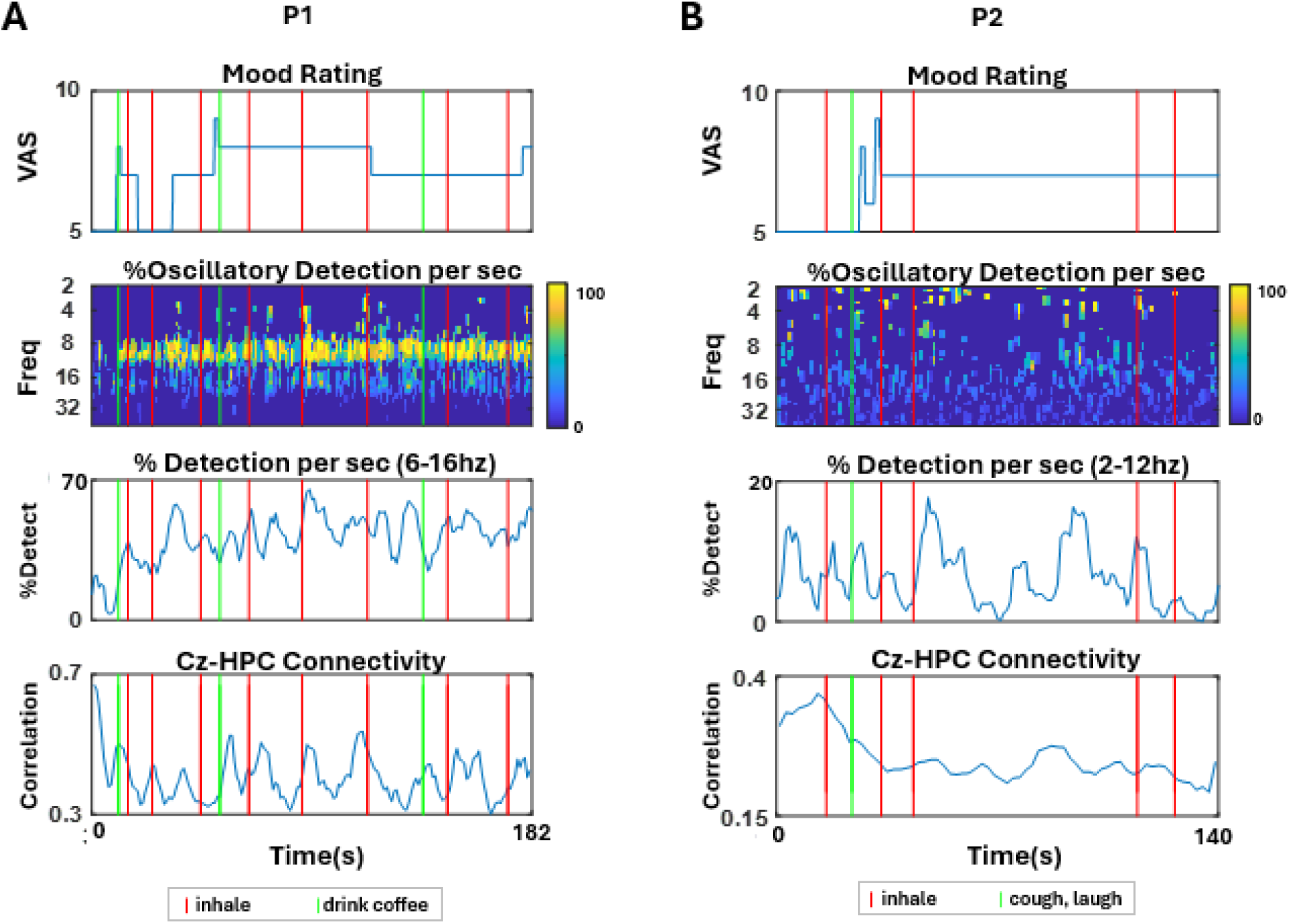
Representative time series of VAS mood, oscillatory detection, and hippocampal-cortical connectivity for P1 (A) and P2 (B).

### Neural data acquisition

For both participants, the RNS electrode leads (NeuroPace) were implanted bilaterally to operate in a closed-loop manner, delivering electrical stimulation to normalize brain activity prior to seizure onset. To avoid stimulation artifacts in the recorded brain activity, stimulation was temporarily disabled during the task with the participants’ informed consent, and the RNS leads were used solely to record intracranial EEG (iEEG) activity. The recording amplifier settings were modified from the clinical default configuration to use a 1 Hz high-pass filter and a 90 Hz low-pass filter, with signals sampled at 250 Hz. Each depth electrode lead contained four contacts with 3.5 mm or 10mm spacing between adjacent contacts, from which local field potentials (LFP) were recorded. In addition to intracranial recordings, scalp EEG was recorded using a WaveGuard 64-channel cap (ANT Neuro, Hengelo, The Netherlands) arranged in an equidistant montage, with the reference at 5Z and the ground at 0Z according to the manufacturer’s numeric electrode labeling scheme).

iEEG and scalp EEG signals were synchronized using a custom-built wand tool that injected a marking artifact into the real-time iEEG and scalp EEG recordings, allowing the two signals to be aligned during analysis [16].

### Neural data preprocessing

To determine the precise anatomical location of each electrode contact from the RNS leads, postoperative high-resolution head computed tomography (CT) images were co-registered with preoperative high-resolution structural magnetic resonance imaging (MRI; T1- and/or T2-weighted sequences) for both participants. Recording contacts were configured in a bipolar montage and re-referenced. Contacts located in the hippocampus with highest signal to noise ratio were selected for subsequent analyses. For P1, contacts 3–4 on the left lead and contacts 1-2 on the right lead were used. For P2, all contacts on the left lead were excluded due to excessive noise, and only contacts 1-2 on the right lead were included in the analysis. Bipolar-referenced LFP signals from these selected contacts were used for subsequent analyses.

Scalp EEG data were preprocessed using the PREP pipeline [17] and EEGLAB [18] toolboxes in MATLAB. Channels corresponding to the standard 10-10 electrode locations Fz, Cz, and Pz were identified from the manufacturer’s numeric electrode labels (2Z, 4Z, and 6Z, respectively) and used for subsequent analyses.

### Frequency of interest

We applied oscillatory analyses focusing on two frequency ranges with complementary rationales. First, we examined the canonical alpha band (8-10 Hz), which has been implicated in affective processing [19,20], cognitive control [21–23], and substance-related behaviors [24,25], particularly within medial frontal and limbic circuits. Given the involvement of these systems in craving and emotional regulation, alpha activity provides a well-motivated band for investigating neural dynamics during nicotine use. Second, we analyzed an individualized frequency band centered on the dominant oscillatory peak observed in the power spectrum. In this dataset, P1 showed dominant power in the 6-16 Hz range, whereas P2 showed dominant power in the 2-12 Hz range, as shown in Figure S1. Because electrophysiological oscillations can vary across individuals and recording sites, focusing on the empirically dominant oscillatory component allows the analysis to capture subject-specific rhythmic activity that may not align precisely with canonical band boundaries. Together, these complementary approaches enable a theory-driven examination of established oscillatory mechanisms alongside a data-driven characterization of the dominant neural dynamics.

### Oscillatory detection and mood

Time-frequency analyses of intracranial EEG (iEEG) and scalp EEG data were performed using the BOSC toolbox [26] to quantify oscillatory activity. Signals were decomposed using a wavelet with a wavenumber of 6 across frequencies spanning 0.71-76.1 Hz, sampled at 60 logarithmically spaced frequency steps.

Oscillatory detection was implemented using the BOSC framework. For each frequency, BOSC estimated the background power spectrum and defined a frequency-specific power threshold as the 95th percentile of the background distribution, along with a duration threshold of 3 cycles. Oscillatory episodes were identified only when power exceeded the threshold for at least the required duration, producing a binary time-frequency detection matrix. For band-specific analyses, detection values were averaged across frequency bins within the band of interest. Detection values were then averaged within 1-s windows to generate a detection time series, reflecting the proportion of each second occupied by detected oscillatory activity (Figure 2).

For the purposes of this study, we focused on the alpha band, given its involvement in reward and pleasure processing. The alpha-band power time series and the VAS mood time series were down-sampled to 1 Hz and smoothed using a 5-point moving average. The two time series were then compared using the Kendall rank correlation coefficient (Kendall’s τ) to assess the ordinal association between these paired observations. This analysis was performed separately for each participant and recording channel. These within-subject correlations were used to characterize subject-specific relationships between alpha activity and mood ratings. We also analyzed individualized frequency bands (6-16 Hz for P1, 2-12 Hz for P2), correlating their oscillatory detection time series with VAS mood time series. Because the analyses were exploratory and interpreted at the individual level rather than for group-level inference, results are presented both with and without correction for multiple comparisons.

In addition, given that the theta band is commonly examined in the context of nicotine and other substance use, particularly in relation to craving [27–29], we performed the same analyses using canonical frequency bands, including the theta band and a combined theta-alpha band, shown in Figure S2.

### Enveloped correlation and nicotine use

Functional connectivity between scalp EEG (Fz, Cz, Pz) and iEEG (hippocampus) signals was assessed using amplitude envelope correlation. Signals were first bandpass filtered using a second-order Butterworth filter to isolate activity within the frequency range of interest. The analytic amplitude envelope of each filtered signal was then extracted using the Hilbert transform. To ensure temporal alignment between recording systems, scalp EEG signals were downsampled to 250 Hz to match the iEEG sampling rate.

Cross-correlation of the amplitude envelopes was then quantified using a sliding-window approach. For P1, where the dominant frequency band was 6-16 Hz, correlations were computed within 4-second windows with 80% overlap between consecutive windows. For P2, whose dominant frequency band was 2-12 Hz, correlations were computed within 10-second windows with 80% overlap between windows. For analyses limited to the alpha band (8-10 Hz), 3-second windows with no overlap were used. Window lengths were selected to capture approximately 20 cycles of the lowest frequency within each band. The degree of overlap was chosen to maximize the number of analyzable windows within the available recording duration, with larger overlaps used for longer window lengths.

For each window, cross-correlation was evaluated across temporal lags up to ±3 seconds, and the peak correlation value was extracted to characterize the strength of envelope coupling between the two signals. This procedure produced a time series reflecting the evolving envelope-based connectivity between the scalp EEG and iEEG channels.

To examine how functional connectivity changed with mood, the connectivity time series was downsampled to 1 Hz and smoothed using a 5-point moving average. For the alpha band connectivity, the alpha detection time series was correlated with the VAS mood time series using the Kendall rank correlation coefficient (Kendall’s τ). This analysis was performed separately for each recording channel pair. For the participant-specific frequency bands, statistical significance was assessed using a circular-shift permutation test based on Kendall’s τ, because the connectivity time series were derived from overlapping windows and therefore contained temporally non-independent samples. In this test, the observed Kendall correlation was first computed between the paired time series. A null distribution was then generated by circularly shifting one time series relative to the other and recomputing Kendall’s τ across 10,000 permutations, with shifts constrained to be at least 5 samples to avoid trivial offsets. The empirical two-sided p-value was defined as the proportion of permuted τ values whose magnitude exceeded or equaled the observed τ. For both statistical tests, false discovery rate (FDR) correction [30] was applied across the three channel pairs to account for multiple comparisons.

To examine broader changes in connectivity beyond moment-to-moment fluctuations, we also compared connectivity during early versus later times of smoking. Specifically, we extracted the 30 seconds preceding the second inhalation and compared it with the final 30 seconds of the recording. Because limited data were available prior to the first inhalation (particularly for P1), we selected the 30-second segment before the second inhalation to represent an early phase of smoking that still reflected the initial mood changes associated with nicotine exposure. To maintain independence among data points within each 30-second segment, consecutive windows were extracted without overlap. As a result, the number of observations per segment was small (N < 10), limiting the statistical power of conventional significance tests. Therefore, we computed Cliff’s delta (δ), a nonparametric effect size measure, to characterize the direction and magnitude of connectivity changes associated with nicotine use.

## Results

### Changes in craving following nicotine use

Both participants completed the Questionnaire of Smoking Urges (QSU) to quantify nicotine craving before and after nicotine use (Table 2). Overall craving scores were higher in P1, who did not have depression but had greater nicotine dependence (per FTND score; see Table 1), compared with P2, who had depression. In both participants, appetitive craving scores were higher than relief craving scores. Following nicotine use, P1’s craving scores decreased to the minimum value (score = 1), indicating the absence of an urge to smoke. In contrast, P2 showed only minimal reductions in craving after nicotine use. This difference was most apparent in the relief craving component, which remained the same for P2.

**Table 2.** Questionnaire of Smoking Urges (QSU) scores before and after nicotine use.

| Participant ID | P1 | P2 |
| --- | --- | --- |
| **QSU before nicotine use** |  |  |
| Appetitive craving | 6.4 | 6.1 |
| Relief craving | 3.1 | 2.2 |
| **QSU after nicotine use** |  |  |
| Appetitive craving | 1 | 5.8 |
| Relief craving | 1 | 2.2 |

QSU scores represent mean ratings (1-7) within each factor, with higher values indicating greater craving.

### Oscillatory activity tracks mood during nicotine use

While smoking/vaping, participants continuously rated their mood, allowing us to examine the moment-to-moment relationship between mood and neural oscillatory activity. Mood ratings ranged from 5 (neutral) to 8 (with 10 indicating very happy) for both participants. Oscillatory detections were correlated with mood using both participant-specific dominant frequency bands (P1: 6-16 Hz; P2: 2-12 Hz) and the canonical alpha band (8-10 Hz). (Figure 3)

**Figure 3.**
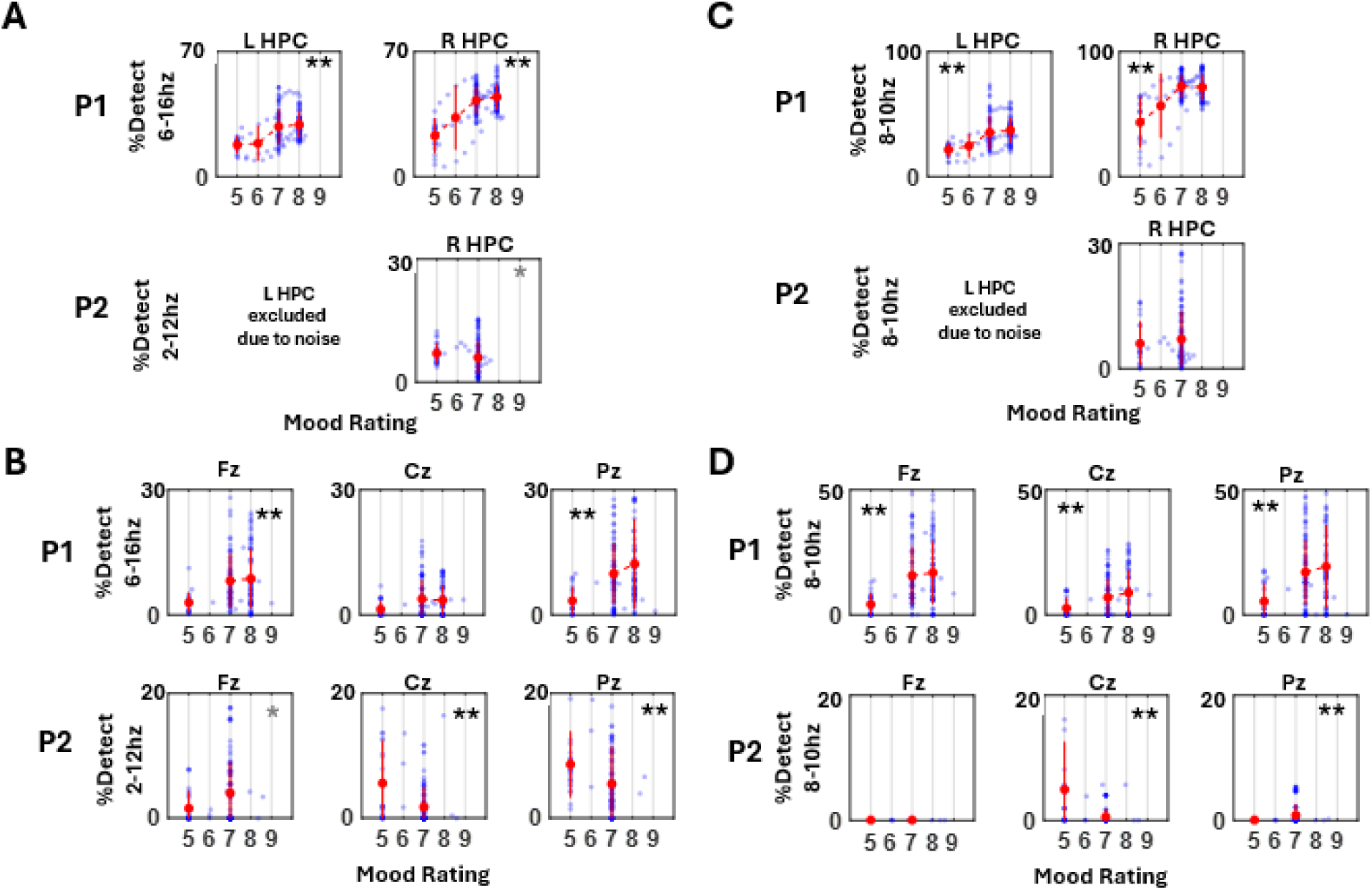
Oscillatory activity tracks mood in individualized frequency bands (A-B) and the alpha band (C-D) for the hippocampus (A, C) and scalp EEG (B, D). HPC = hippocampus. Black asterisks indicate statistical significance after FDR correction; gray asterisks indicate significance without FDR correction. * indicates *p* < 0.05; ** indicates *p* < 0.01.

For P1, mood ratings showed a consistent positive association with oscillatory detection across both the personalized and alpha frequency bands. This relationship was observed in bilateral hippocampal recordings as well as scalp electrodes (Fz, Cz, Pz).

Kendell correlation coefficients ranged from approximately τ = 0.15 to 0.31 across regions. For P2, the relationship between mood and oscillatory detection was more mixed. In the hippocampus, the personalized frequency band showed a modest negative correlation with mood, while no association was observed in the alpha band. Across scalp electrodes, the personalized band showed a positive correlation at Fz but negative correlations at Cz and Pz. Within the alpha band, oscillatory activity was not detected at Fz, while correlations were negative at Cz and positive at Pz. Full correlation statistics for each channel are reported in Table S1.

### Reduced hippocampal-cortical connectivity during nicotine use

Using envelope cross-correlation, we examined limbic-cortical functional connectivity between the hippocampus and midline scalp electrodes (Fz, Cz, and Pz). We first focused on the alpha band and examined how this connectivity measure related to mood during real-time nicotine inhalation (Figure 4). In P1, connectivity between the left hippocampus and cortical electrodes showed little association with mood. In contrast, connectivity between the right hippocampus and midline cortical regions showed a negative relationship with mood, such that connectivity decreased as mood ratings increased. A similar pattern was observed in P2, although the associations were more modest.

**Figure 4.**
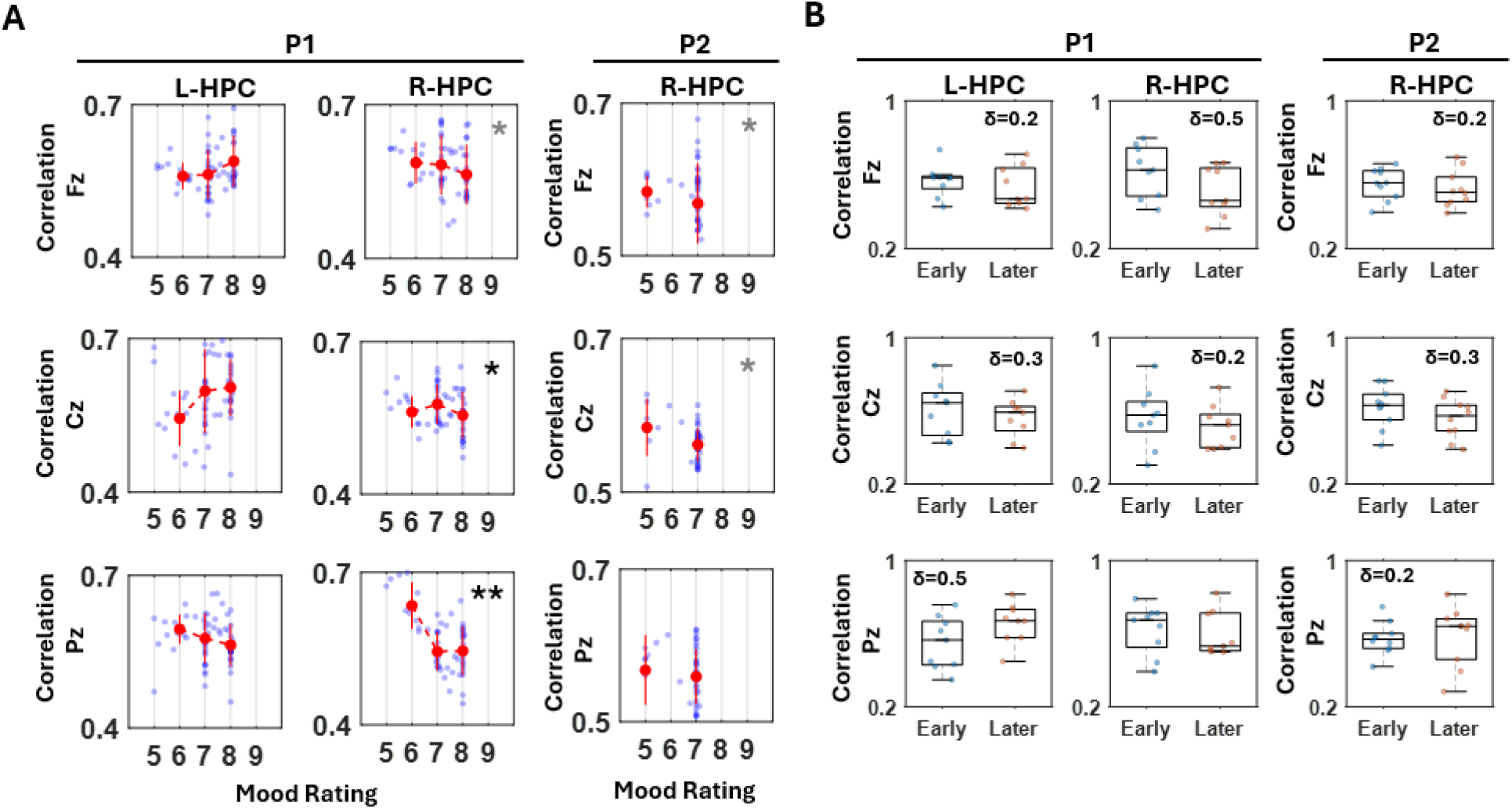
(A) Within the alpha band, hippocampal-cortical connectivity shows a modest negative correlation with mood elevation. (B) Connectivity decreases from early to later stages of nicotine use, particularly at Fz and Cz. Cliff’s delta value is omitted for negligible effect sizes (δ < 0.15) [31]. HPC = hippocampus. Black asterisks indicate statistical significance after FDR correction; gray asterisks indicate uncorrected significance. * indicates *p* < 0.05; ** indicates *p* < 0.01.

We next compared connectivity during later stages of nicotine use with earlier stages, given that overall mood improved following nicotine inhalation in both participants.

Consistent with the moment-to-moment findings, connectivity generally decreased during the later phase of nicotine use. This reduction was most consistently observed for hippocampal connections with Fz and Cz in both participants. In contrast, connectivity involving Pz showed mixed patterns, with increases observed for the left hippocampus in P1 and the right hippocampus in P2.

Using the same analytic pipeline, we also examined connectivity within the participant-specific dominant frequency bands (P1: 6-16 Hz; P2: 2-12 Hz) (Figure 5). Within these personalized bands, the relationship between connectivity and mood showed mixed patterns. For P1, connectivity across regions showed little association with mood changes. In contrast, for P2, the pattern resembled the alpha-band findings, with connectivity decreasing as mood ratings increased. We also compared connectivity during earlier versus later periods of smoking. For P1, only negligible to small reductions in connectivity were observed. In contrast, P2 showed larger reductions in connectivity during the later phase of nicotine use, with the most pronounced effect observed at Cz. Full connectivity statistics for each channel are reported in Table S2.

**Figure 5.**
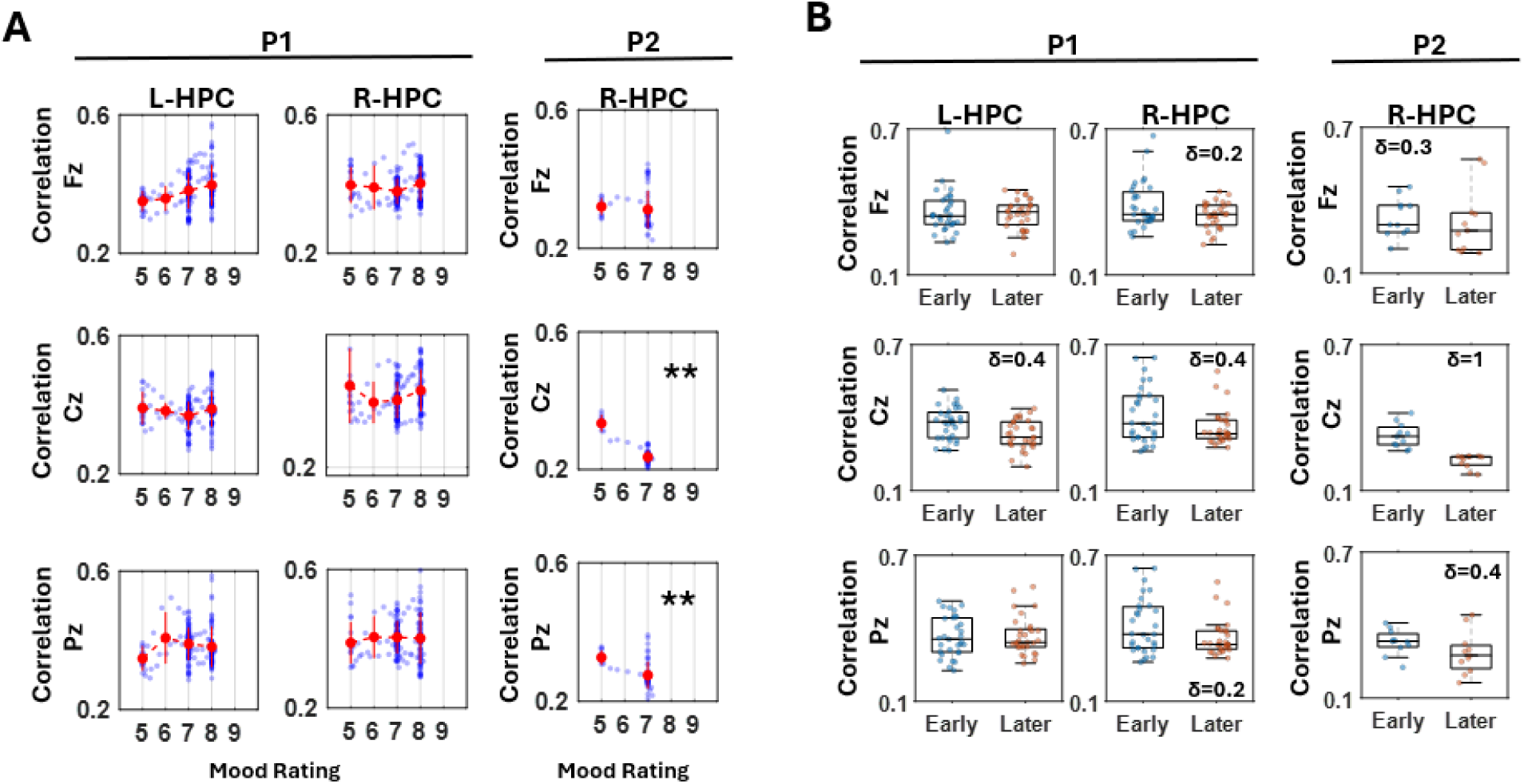
(A) Within the participant-specific dominant frequency band, hippocampal-cortical connectivity shows a negative correlation with mood elevation for P2. (B) Connectivity decreases from early to later stages of nicotine use, more prominently for P2. Cliff’s delta value is omitted for negligible effect sizes (δ < 0.15) [31]. HPC = hippocampus. Black asterisks indicate statistical significance after FDR correction; gray asterisks indicate uncorrected significance. * indicates p < 0.05; ** indicates p < 0.01

## Discussion

This study provides a rare opportunity to examine real-time neural dynamics during nicotine inhalation using simultaneous hippocampal and scalp EEG recordings. This approach enables characterization of limbic-cortical coupling during acute nicotine use and its modulation by depressive status. Although based on two cases, these findings provide hypothesis-generating evidence that depressive status may influence reward-related neural dynamics, motivating further investigation in larger cohorts.

### Oscillatory activity and nicotine use

Oscillatory detection was overall more prevalent in P1 (the participant without depressive symptoms) compared with P2. The dominant oscillatory bandwidth also differed between participants: P1 showed prominent activity in the 6-16 Hz range, whereas P2 showed a broader lower-frequency range of 2-12 Hz. In P1, oscillatory detection increased as mood ratings increased, whereas P2 showed either no association or a modest decrease with increasing mood. These patterns were observed both within the participant-specific dominant frequency bands and within the canonical alpha band (8-10 Hz). Similar results were also observed when using band power (rather than BOSC detection), indicating a consistent overall pattern (Figure S1).

These differences in baseline oscillatory profiles and their relationship to mood may be related to depressive status. The weaker or absent positive association between oscillatory activity and mood in P2 may reflect a diminished rewarding response to nicotine. This interpretation is consistent with the behavioral findings, where P1 showed a marked reduction in craving following nicotine use, whereas P2 exhibited only modest reductions. Together, these results suggest that depressive status may influence reward- and craving-related neural dynamics during nicotine use.

Notably, these patterns were generally consistent across other frequency ranges (4-12 Hz, theta 4-8 Hz, and beta 12-20 Hz), suggesting that the effect was not strongly band-specific (Figure S2). Prior studies have often reported decreases in lower-frequency power (e.g., theta or delta) and increases in higher-frequency power during nicotine use and the opposite during craving [32–34]. However, given the frequency-wide pattern observed here, one possible explanation for this discrepancy is that prior studies may have included heterogeneous participant groups in which psychiatric status was not considered. Future studies with larger samples will be needed to determine whether nicotine-related oscillatory dynamics differ systematically as a function of psychiatric conditions such as depression.

### Reduced hippocampal-cortical connectivity after nicotine use is associated with a shift from reward-seeking to reward and relief processing

Given prior reports of reduced hippocampal and frontoparietal connectivity in depression [35,36], we hypothesized that nicotine use would increase hippocampal-prefrontal connectivity, potentially reflecting a normalization of reward- and relief-related processing, with a greater effect in the participant with depression. However, we observed the opposite pattern. Hippocampal-cortical connectivity decreased following nicotine use and the associated elevation in mood, most prominently in the 8-10 Hz range between the hippocampus and Fz/Cz.

One possible interpretation is that low-alpha activity, which has been linked to behavioral inhibition and regulatory control [21–23], diminishes once the reward from nicotine use is achieved. In this state of satiety, reduced connectivity may reflect a transition from reward-seeking to reward/relief related processing - consistent with the subjective experience of “taking the edge off.” This aligns with prior findings showing stronger prefrontal coupling during drug cue exposure and craving [37–39], suggesting that such connectivity may be stronger during anticipatory or reward-seeking states rather than after reward attainment. The findings with Cz, often associated with motor readiness [40], may further support this interpretation. During craving or preoccupation with obtaining nicotine, the body may be in a heightened state of motor preparedness. The observed reduction in connectivity may therefore reflect a release from this preparatory tension once the reward is attained.

Within a network-based framework of affect [41], these findings suggest that reward and relief are not localized to a single region but instead emerge from interactions among systems integrating cognitive, somatic, and affective processes. In this view, cognitive systems support evaluation and planning for reward acquisition, somatic systems reflect bodily readiness, and posterior midline activity (approximated by Pz) may index internally directed or interoceptive states associated with reward processing. Low-alpha activity, often linked to behavioral inhibition and regulatory control [21–23], may therefore reflect the degree of coordinated engagement across these systems during reward-seeking states such as craving. From this perspective, the observed reduction in low-alpha connectivity following nicotine use may reflect a shift from coordinated, goal-directed reward-seeking to a more settled, less effortful state following relief of craving (Figure 6).

**Figure 6.**
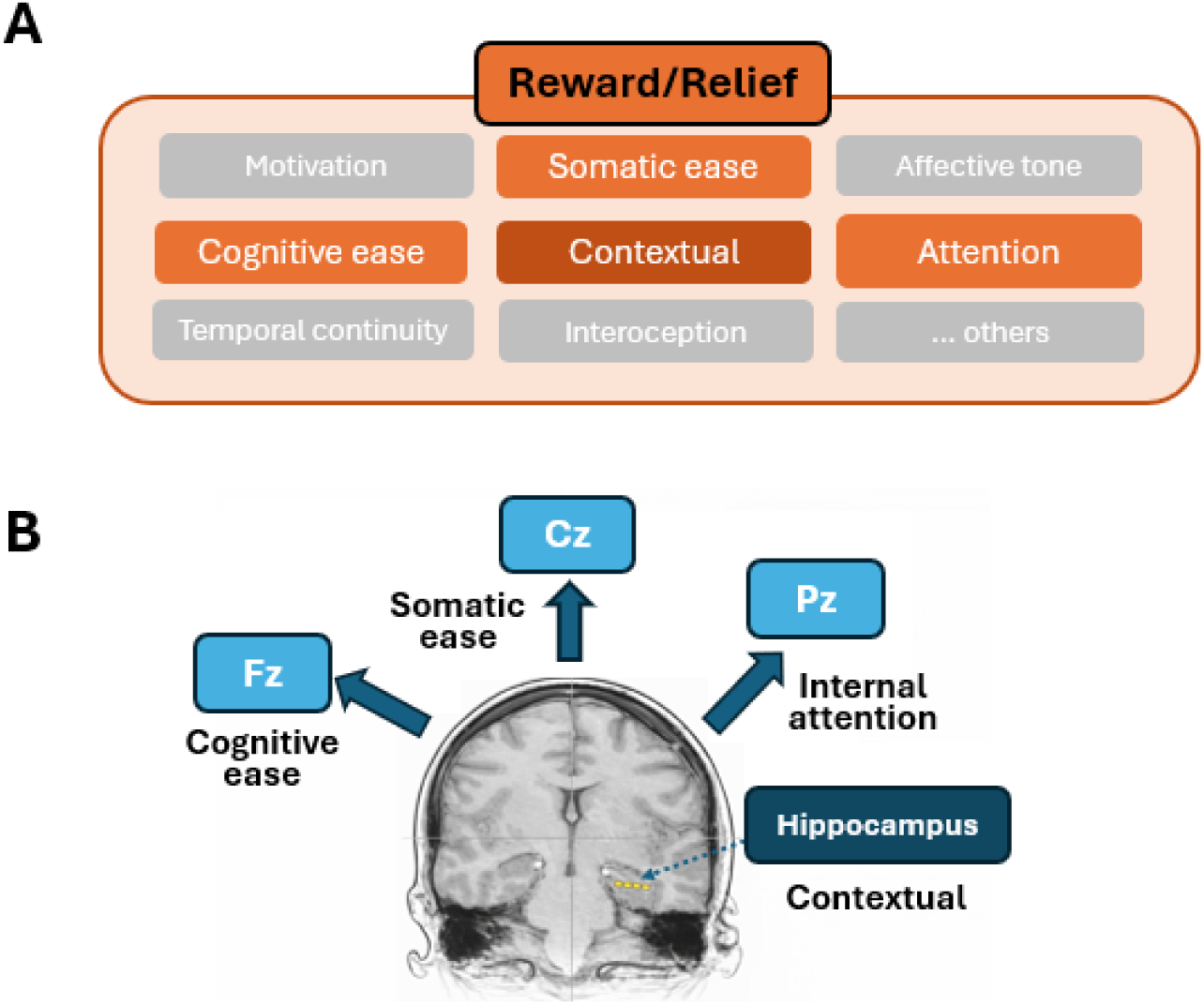
Conceptual framework of network-level processes underlying reward and relief. (A) Schematic illustration of cognitive, somatic, contextual, and affective factors that contribute to the subjective experience of reward and relief. (B) Recording sites used to capture these processes, including hippocampal (contextual) and midline scalp EEG regions (Fz: cognitive processing; Cz: somatic processing; Pz: internally directed attention), providing a systems-level view of limbic-cortical interactions.

### Neural subtypes inform personalized treatment

Heterogeneity among nicotine users is well recognized, but treatment approaches remain largely uniform. In this study, we aimed to examine how acute nicotine use differs between two individuals-one without depression and one with depression-with the ultimate goal of informing more personalized treatment strategies.

Here, we found that oscillatory activity differed markedly between P1 and P2, particularly in relation to mood changes. In P1, oscillatory activity increased with mood elevation across all recorded structures, whereas P2 showed little to no relationship-or even a negative association-between oscillatory activity and mood. In contrast, both participants exhibited reduced hippocampal-cortical connectivity during nicotine use. In addition, P2 reported minimal reduction in relief-related craving and only modest decrease in appetitive craving. This pattern suggests that oscillatory activity may be more closely related to relief craving, whereas reductions in hippocampal-cortical connectivity may reflect appetitive reward or pleasure. However, given the small sample size, these interpretations should be considered preliminary.

Nonetheless, these findings highlight distinct neural dynamics across individuals and provide an initial step toward identifying meaningful subtypes of nicotine users, which may be critical for developing personalized treatments. For example, in alcohol use disorder, reward- and relief-driven subtypes have been distinguished, with reward-driven individuals showing greater benefit from opioid antagonists such as naltrexone [42,43]. Analogously, identifying subtypes in nicotine use may inform targeted interventions, including tailoring repetitive transcranial magnetic stimulation (rTMS) targets and stimulation parameters to individual neural profiles [44–46]. At present, there is limited understanding of how neuromodulatory treatments differentially affect patient subtypes, particularly in the presence of comorbid conditions such as depression. This study therefore provides a starting point for investigating how such variability shapes nicotine-related neural responses and informs treatment strategies.

### Limitation

Although this study provides a useful starting point, several limitations should be considered so that future research can address them in larger cohorts. First, including appropriate control conditions will be critical to account for potential confounding factors. For example, incorporating a condition in which participants perform a similar motor action-such as reaching for and drinking water-would help control for action-related effects. Additionally, including other naturalistic pleasurable activities that are independent of substance use (e.g., watching a humorous video) could help disentangle the effects of nicotine intoxication from general reward-related responses.

Another limitation is the minimal control over participants’ prior nicotine use. Because recordings were conducted opportunistically during breaks between other research studies, we were unable to standardize abstinence periods (e.g., requiring participants to refrain from nicotine use for a set number of hours prior to recording). As a result, differences observed between participants (e.g., P1 and P2) cannot be attributed solely to variables of interest, such as the presence of depression. Other factors-including recent nicotine use, age, sex, anxiety diagnosis, method of nicotine delivery (vaping vs. cigarettes), and level of nicotine dependence (see Table 1)-may have contributed to the observed differences.

Furthermore, the limited number of puff instances per participant constrains the robustness of the findings. Future studies should include a larger dataset collected across multiple sessions per participant and, ideally, span several days in naturalistic settings. Such an approach would allow for better control of both within- and between-subject variability and provide a more reliable assessment of nicotine-related effects.

## Conclusion

In summary, this study demonstrates capturing real-time limbic-cortical dynamics during nicotine use using simultaneous hippocampal and scalp EEG recordings. The findings reveal distinct neural patterns across individuals, suggesting that depressive status may influence reward- and relief-related processes. Oscillatory activity and hippocampal-cortical connectivity appear to reflect different aspects of nicotine-related states, supporting a network-based framework in which reward and relief emerge from distributed interactions. Although preliminary, these results highlight the potential for identifying neurophysiological subtypes of nicotine users and provide a foundation for developing personalized treatment approaches in future studies.

## Acknowledgement

This research was supported by an NIH K01 Mentored Research Scientist Development Award (1K01DA060327) for JR. We thank Nanthia Suthana for the support and feedback on this work.

## Data and code availability

The data and custom computer code used to generate results are available from the corresponding author upon request.

## Supplemental Materials

**Table S1.** Statistical results for correlations between oscillatory activity and mood.

| Participant ID | P1 | P2 |
| --- | --- | --- |
| **Fig 3A-B Kendall Tau** |  |  |
| L-HPC | 0.26 (p<0.001) | n/a |
| R-HPC | 0.27 (p<0.001) | -0.16 (p=0.02) |
| Fz | 0.17 (p=0.002) | 0.16 (p=0.03) |
| Cz | 0.10 (p=0.09) | -0.26 (p<0.001) |
| Pz | 0.16 (p=0.0051) | -0.22 (p<0.001) |
| **Fig 3C-D Kendall Tau** |  |  |
| L-HPC | 0.31 (p<0.001) | n/a |
| R-HPC | 0.17 (p=0.0016) | -0.01 (p=0.84) |
| Fz | 0.22 (p<0.001) | nan |
| Cz | 0.15 (p=0.010) | -0.37 (p<0.001) |
| Pz | 0.15 (p=0.006) | 0.29 (p<0.001) |
| **Fig S1B-C Kendall Tau** |  |  |
| L-HPC | 0.14 (p=0.01) | n/a |
| R-HPC | 0.39 (p<0.001) | -0.17 (p=0.013) |
| Fz | -0.06 (p=0.32) | -0.44 (p<0.001) |
| Cz | -0.08 (p=0.13) | -0.23 (p<0.001) |
| Pz | 0.14 (p=0.01) | -0.39 (p<0.001) |
| **Fig S2A-B Kendall Tau** |  |  |
| L-HPC | 0.32 (p<0.001) | n/a |
| R-HPC | 0.25 (p<0.001) | 0.12 (p=0.07) |
| Fz | 0.19 (p=0.001) | 0.07 (p=0.35) |
| Cz | 0.09 (p=0.14) | -0.25 (p=0.001) |
| Pz | 0.16 (p=0.004) | -0.008 (p=0.91) |
| **Fig S2C-D Kendall Tau** |  |  |
| L-HPC | 0.31 (p<0.001) | n/a |
| R-HPC | 0.25 (p<0.001) | -0.26 (p<0.001) |
| Fz | 0.15 (p=0.007) | 0.07 (p=0.35) |
| Cz | 0.01 (p=0.86) | -0.21 (p=0.005) |
| Pz | 0.13 (p=0.03) | -0.03 (p=0.69) |
| **Fig S2E-F Kendall Tau** |  |  |
| L-HPC | 0.15 (p=0.007) | n/a |
| R-HPC | 0.34 (p<0.001) | 0.01 (p=0.92) |
| **Fz** | -0.02 (p=0.68) | nan |
| **Cz** | 0.04 (p=0.51) | -0.08 (p=0.28) |
| **Pz** | 0.12 (p=0.03) | -0.04 (p=0.56) |

**Table S2.** Statistical results for hippocampal–cortical connectivity analyses.

| **Participant ID** | **P1**<br>**L-HPC** | **P1**<br>**R-HPC** | **P2**<br>**R-HPC** |
| --- | --- | --- | --- |
| **Fig 4A Kendall tau** |  |  |  |
| **Fz** | 0.13 (p=0.2) | -0.20 (p=0.03) | -0.26 (p=0.02) |
| **Cz** | 0.16 (p=0.08) | -0.24 (p=0.01) * | -0.24 (p=0.04) |
| **Pz** | -0.15 (p=0.1) | -0.37 (p<0.01) ** | -0.10 (p=0.4) |
| **Fig 4B Cliff Delta** |  |  |  |
| **Fz** | -0.23 | -0.46 ^ | -0.20 |
| **Cz** | -0.28 | -0.19 | -0.30 |
| **Pz** | 0.46 ^ | -0.14 | -0.22 |
| **Fig 5A Kendall tau** |  |  |  |
| **Fz** | 0.17 (p=0.2) | 0.10 (p=0.3) | -0.20 (p=0.3) |
| **Cz** | 0.06 (p=0.6) | 0.11 (p=0.2) | -0.52 (p<0.01) ** |
| **Pz** | -0.02 (p=0.9) | -0.02 (p=0.8) | -0.42 (p<0.01) ** |
| **Fig 5B Cliff Delta** |  |  |  |
| **Fz** | 0.05 | -0.17 | -0.26 |
| **Cz** | -0.38 ^ | -0.35 ^ | -1.0 ^^ |
| **Pz** | 0.01 | -0.24 | -0.44 ^ |

**Figure S1.**
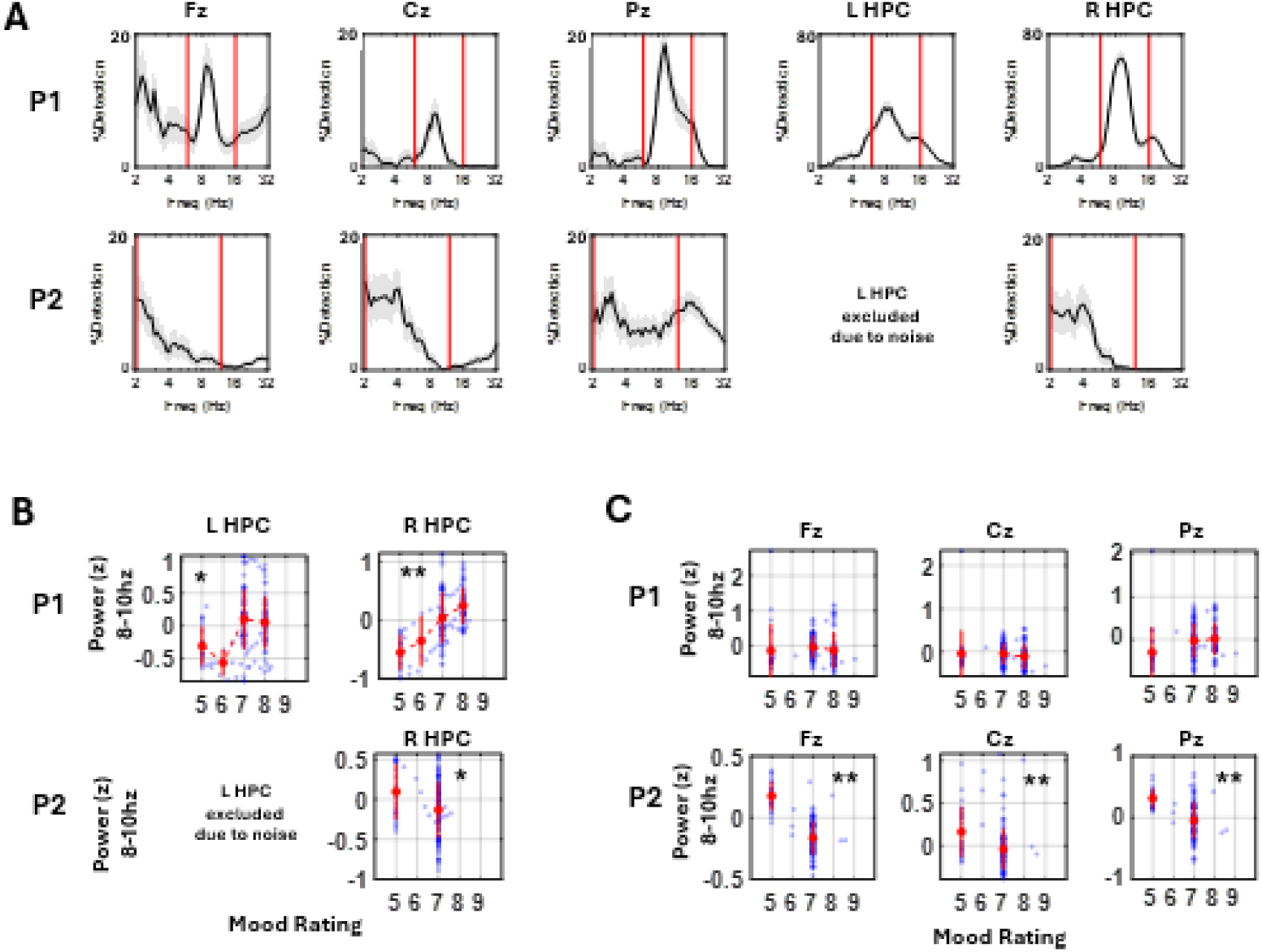
Oscillatory activity across recording structures. (A) Oscillatory detection rate across frequency 2-32hz. (B) Oscillatory activity quantified as band power (z-scored across the recording duration) and its correlation with mood ratings. HPC = hippocampus. Black asterisks indicate statistical significance after FDR correction. * indicates *p* < 0.05; ** indicates *p* < 0.01.

**Figure S2.**
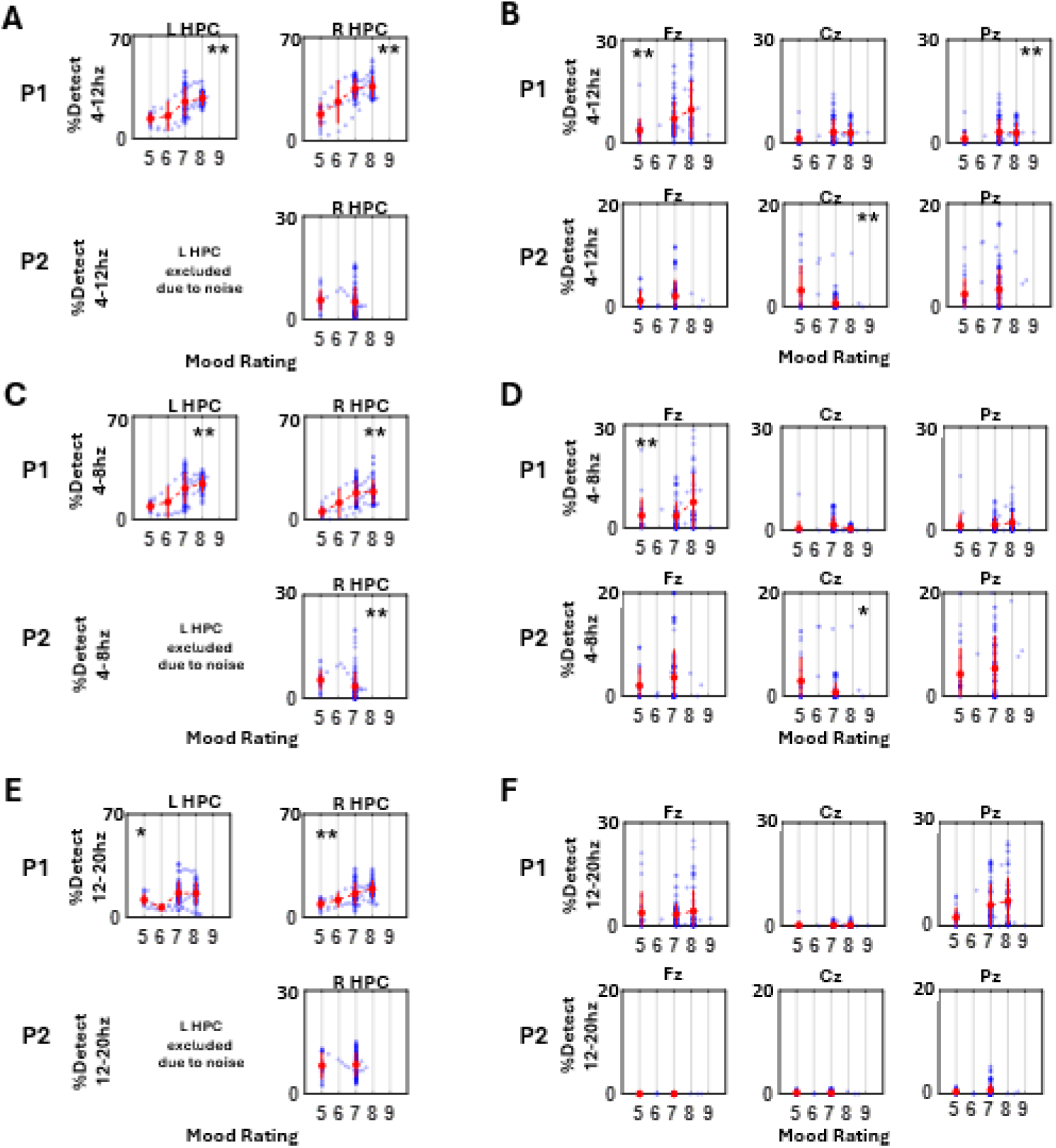
Oscillatory activity and mood across frequency bands. Theta–alpha (A, B), theta (C, D), and beta (E, F) bands are shown for the hippocampus and scalp EEG. HPC = hippocampus. Black asterisks indicate statistical significance after FDR correction; gray asterisks indicate uncorrected significance. * indicates p < 0.05; ** indicates p < 0.01.

